# Cnidarian cell type diversity revealed by whole-organism single-cell RNA-seq analysis

**DOI:** 10.1101/201103

**Authors:** Arnau Sebé-Pedrós, Elad Chomsky, Baptiste Saudememont, Marie-Pierre Mailhe, Flora Pleisser, Justine Renno, Yann Loe-Mie, Aviezer Lifshitz, Zohar Mukamel, Sandrine Schmutz, Sophie Nouvault, Francois Spitz, Amos Tanay, Heather Marlow

## Abstract

A hallmark of animal evolution is the emergence and diversification of cell type-specific transcriptional states. But systematic and unbiased characterization of differentiated gene regulatory programs was so far limited to specific tissues in a few model species. Here, we perform whole-organism single cell transcriptomics to map cell types in the cnidarian *Nematostella vectensis*, a non-bilaterian animal that display complex tissue-level bodyplan organization. We uncover high diversity of transcriptional states in *Nematostella*, demonstrating cell type-specific expression for 35% of the genes and 51% of the transcription factors (TFs) detected. We identify eight broad cell clusters corresponding to cell classes such as neurons, muscles, cnidocytes, or digestive cells. These clusters comprise multiple cell modules expressing diverse and specific markers, uncovering in particular a rich repertoire of cells associated with neuronal markers. TF expression and sequence analysis defines the combinatorial code that underlies this cell-specific expression. It also reveals the existence of a complex regulatory lexicon of TF binding motifs encoded at both enhancer and promoters of *Nematostella* tissue-specific genes. Whole organism single cell RNA-seq is thereby established as a tool for comprehensive study of genome regulation and cell type evolution.

---

Non-bilaterian animal lineages, including cnidarians, ctenophores, sponges and placozoans, have simple body plans and have been historically considered to contain limited numbers of cell types (Valentine, 2003). In cnidarians, these cells include a small number of morphologically distinct neurons, gland cells, muscle cells, epidermis, and gut (Frank and Bleakney, 1976; Hand and Uhlinger, 1992; Hyman, 1940). This presumed simplicity in the number of cnidarian cell types stands in marked contrast to their genomic complexity. Indeed, multiple genomic features, such as the gene repertoire, syntenic gene blocks, and intronic structure, are more similar between cnidarians and vertebrates than to model bilaterian invertebrates like *D. melanogaster* or *C. elegans* (Putnam et al., 2007; Technau and Schwaiger, 2015). Recently, it was shown that these similarities extend to the regulatory landscape of *Nematostella* genes (Schwaiger et al., 2014). However, the extent to which these genomic features participate in regulating complex cell type hierarchies remains largely unknown.

Gene expression profiling by *in situ* hybridization allows for comparative study of cell types and tissue organisation in different species, providing insights into their evolution (Steinmetz et al., 2012). But these approaches require *a priori* selection of candidate gene markers; they are difficult to scale towards multiple expressed genes simultaneously; and they are not readily applicable to all species or life stages, in particular adult specimens. On the other hand, techniques for genome-wide profiling of gene expression were so far dependent on established staging and tissue dissection procedures. Single-cell RNA sequencing (scRNA-seq) is rapidly emerging as a powerful approach for unbiased *de novo* discovery and detailed molecular characterization of transcriptional states and cell types in mammalian tissues (Jaitin et al., 2014; La Manno et al., 2016; Shekhar et al., 2016; Tasic et al., 2016; Wurtzel et al., 2015; Zeisel et al., 2015). Nevertheless, systematic single cell expression profiling of whole adult non-model organisms remains a challenge yet to be explored.

We focused our whole-organism scRNA-seq efforts on the cnidarian *Nematostella* (Rentzsch and Technau, 2016), hoping to rely on the relatively well-established understanding of basic cell types and developmental processes in this species, and to expand them using unbiased and high-resolution analysis. We were particularly interested in the organization of neuronal cell types in *Nematostella*, given the important role of *Nematostella* in evolutionary studies of the nervous system (Marlow et al., 2009; Nakanishi et al., 2012). We applied MARS-seq (Jaitin et al., 2014) to whole adults, using fluorescence-activated cell sorting (FACS) to distribute live cells into 384-well plates and performed labelling, amplification, sequencing and de-noising as previously described (Jaitin et al., 2014). We complemented whole adult data with single cell profiles derived from juvenile polyps as well as specific subsets of both micro-dissected and FACS-sorted cell populations. More than earlier applications of scRNA-seq for cell type dissection or characterization of transcriptional variation within cell types, whole-organism analysis in *Nematostella* involved dramatic variability in cell size and total RNA-content within the data cohort (**Figure 1A**). We therefore lowered the coverage threshold on single cells to 100 molecules, and developed an analytic pipeline that combines non-parametric construction of K-nearest neighbour graphs, detection of cell modules and multiple layers of validation using direct tests for module coherence and bootstrap iterations. This facilitate the dissection a single cell cohort involving up to 100-fold difference in recovered RNA-content. We retained for analysis 11,888 cells that showed sufficiently high QC metrics, including 10,713 adult and 1,175 juvenile polyp cells, with a median number of 541 RNA molecules per cell (**Figure 1A**). Using a combined genome annotation scheme we mapped 81% of the MARS-seq sequences to the genome (**Figure S1A-B**) and detected expression for 16,853 genes (66% of the total predicted genes, with at least 20 total molecules). The global expression distribution (**Figure 1B**) indicates that 17.5% of the genes represented 80% of the transcription, with a heavy tail of low expression transcripts accounting for the remaining 20%.

**Figure 1.**
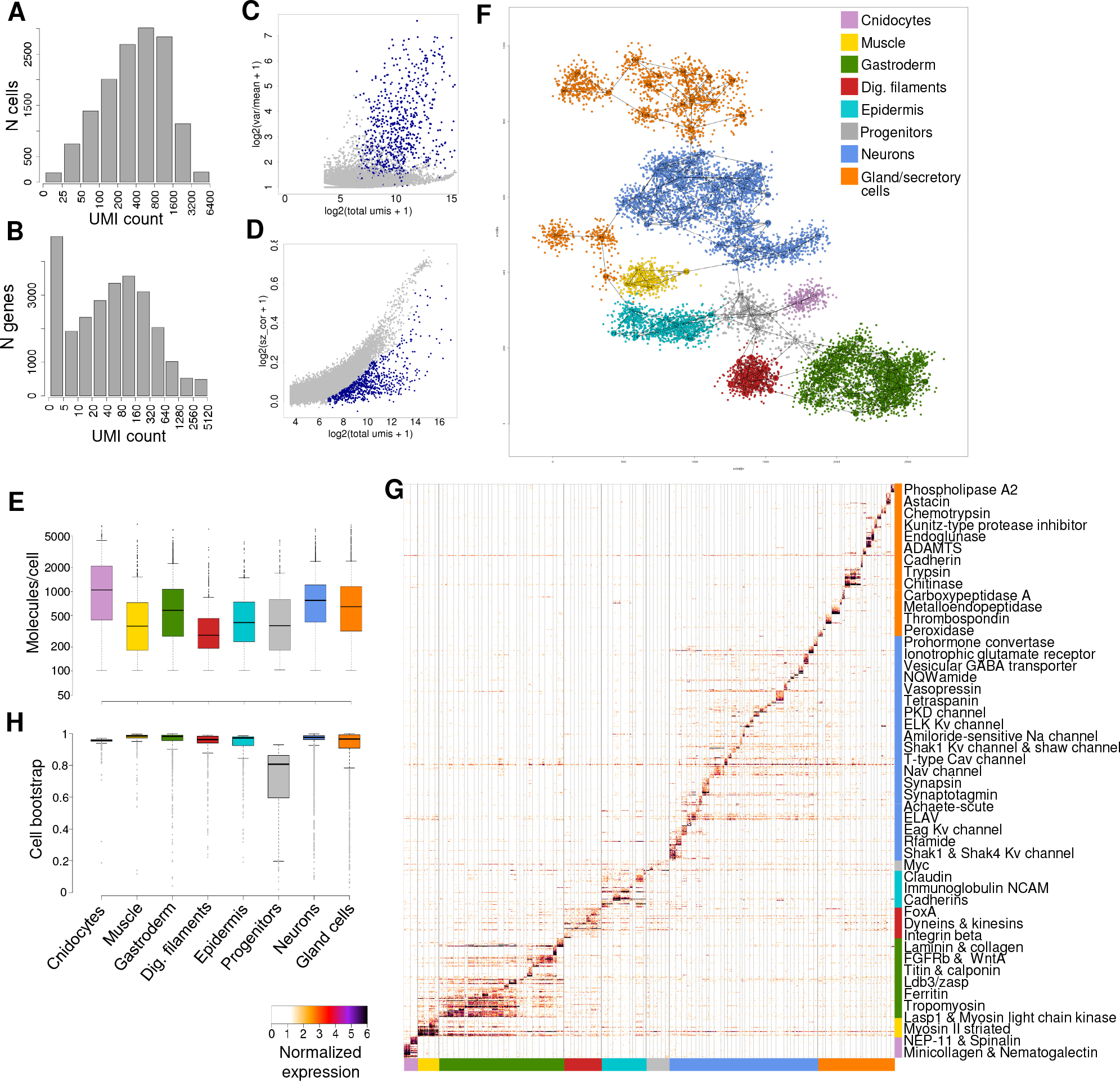
Whole-organism unbiased dissection of *Nematostella* cell types. **(A)** Distribution of total RNA molecules per cell. **(B)** Distribution of total RNA molecules per gene. **(C)** Relationship between gene expression var/mean across cells (y-axis) and gene total expression (x-axis). Marker genes selected for cell clustering are shown in blue. **(D)** Relationship between gene total expression (x-axis) and the correlation between gene expression and total RNA molecules per cell (y-axis). Marker genes selected for cell clustering are shown in blue. **(E)** Distribution of total RNA molecules per cell by broad cluster. **(F)** 2d projection of Nematostella 104 cell modules and 10,460 cells, color-coded according to our broad cluster definitions. **(G)** Normalized expression of 904 genes (rows) across 10,460 cells (columns), sorted by cell module and broad cluster association. For each module, the top 25 genes sorted by fold change versus the other cell modules were selected (with a FC threshold>=2). **(H)** Distribution of broad cell cluster association frequencies in 1,000 bootstrap subsamplings.

To identify genes with potential cell type-specific expression while accounting for variability in cell size, we analysed expression variance across cells (**Figure 1C**), and the correlation of individual gene expression with the total RNA content per cell (**Figures 1D** **and S1C**). We selected 700 genes with high overall expression but low correlation with total RNA-content, providing features that sensitively define the similarity between sparse single cell profiles. We observed 99.5% of the cells are expressing at least seven of the selected markers (**Figure S1D**). Using these gene features, we identified 104 cell modules, and pooled RNA in each module to derive 104 complex transcriptional signatures that are observed in *Nematostella* cells (**Figure S1E-F**). While individual cells in our data may be limited in their mRNA profile depth, modules are pooling together similar cells to allow more quantitative and robust analysis. We performed bootstrap and subsampling analysis to quantify modules robustness (**Figure S1G-H**). This demonstrated that, despite the broad distribution of the number of molecules per single cell in our data (**Figure 1E),** single cells retain high information content and could be grouped robustly into a large number of distinct molecular behaviours. As suggested by a graph based 2D projection plot (**Figure 1F**) and direct visualization of genes distributions over cells (**Figure 1G**), and as further supported by the bootstrap analysis (**Figure 1H**), we organized the complex transcriptional landscape in *Nematostella* into eight broad cell clusters. Based on annotation of enriched marker genes we identified these clusters as cnidocytes, muscle cells, gastroderm, digestive filaments, epidermis, progenitor cells, neurons and gland/secretory cells (**Figure S2**).

The eight broad cell clusters we defined contain additional sub-structure, indicative of transcriptional heterogeneity. But before zooming into these more specific molecular behaviors within the clusters, analysis of the broad cell clusters allows us to globally characterize *Nematostella* cell type hierarchy. In fact, these eight cell groups represent the primary phenotypic cell classes previously described in anthozoan cnidarians such as *Nematostella.* For example, we found that the cnidarian gastrodermal lining and mesenteries, previously termed the “endomesoderm” (Martindale et al., 2004), can be subdivided into two distinct clusters that we term digestive filaments, where prey is held and digested, and the gastrodermal lining. The gastrodermal cell cluster, including the classically described epitheliomuscular cells (Frank and Bleakney, 1976), corresponds to signatures of genes associated with non-fast musculature, including expression of myosin light chain, and several collagens. The digestive filament cells, which represent the functional digestive elements of the animal, including epithelial and cilioglandular structures of the mesenteries, show expression a trp channel, a known mesentery- restricted hedgehog (Matus et al., 2008) and the MPP5 MAGUK protein (**Figure S2**). Cnidocyte cells, which are highly specialized evolutionary novelties that enable prey capture and defense, show co-expression of multiple venom and capsule proteins, such as minicollagens, NEP proteins and nematogalectins, in agreement with the known biology of this cell type (Zenkert et al., 2011) (**Figure S2**). Epidermal cells showed expression of multiple adhesion molecules; while the highly diverse gland/secretory cells were grouped based on their functional similarity, each of them expressing different combinations of proteases and other enzymes, which might be related to digestion but also to non-cnidocyte venom functions (Jaimes-Becerra et al., 2017; Moran et al., 2013). Finally, a group of modules are constituted by putative progenitor cells, which may underlie the regenerative properties of *Nematostella* (Passamaneck and Martindale, 2012). These cells were much less coherently associated than other clusters (**Figure 1H**), but they show expression of several factors associated with stemness (Alié et al., 2015; Hemmrich et al., 2012) (**Figure S2**).

Comparison of the eight *Nematostella* broad cell clusters with published vertebrate organ-specific transcriptomes (Brawand et al., 2011) revealed similarities between this neuronal cluster and vertebrate brain/cerebellum tissues (**Figure S3A)**, as well as linkage between the contractile gastrodermis and muscle *Nematostella* tissues with vertebrate heart transcriptomes. Despite the vast evolutionary timescale separating *Nematostella* from vertebrates, several conserved effector genes, such as ion channels, synaptic components, and the neuron-associated RNA-binding protein ELAV show consistent co-expression that underlie these deep similarities (**Figure S3B)**. We expect richer comparative analyses will be dependent on much denser phylogenetic sampling of non-bilaterian and invertebrate bilaterian species.

In order to position some of our cell modules spatially and validate them against known *Nematostella* morphological features, we performed *in situ* hybridization (ISH) using probes against several genes with restricted, specific expression patterns. We observed remarkable correspondence between the annotation of clusters, the genes associated with them, and the ISH spatial expression maps (**Figure 2A-E**). Markers for specific digestive enzyme-producing gland cells (**Figure 2D**), and muscle cells (**Figure 2B**) showed consistent spatial distributions marking the gland cells of the digestive gut filaments and the longitudinal retractor musculature and “spindle-shaped” tentacle myocytes, respectively (Renfer et al., 2010). Markers for tentacle and body wall epidermal cells showed broad expression in these tissues (**Figure 2C**). In summary, ISH data provide support for the scRNA-based identification of cell types, suggesting that the specific transcriptional signatures observed in modules are consistent with the *Nematostella* spatial cell type organization.

**Figure 2.**
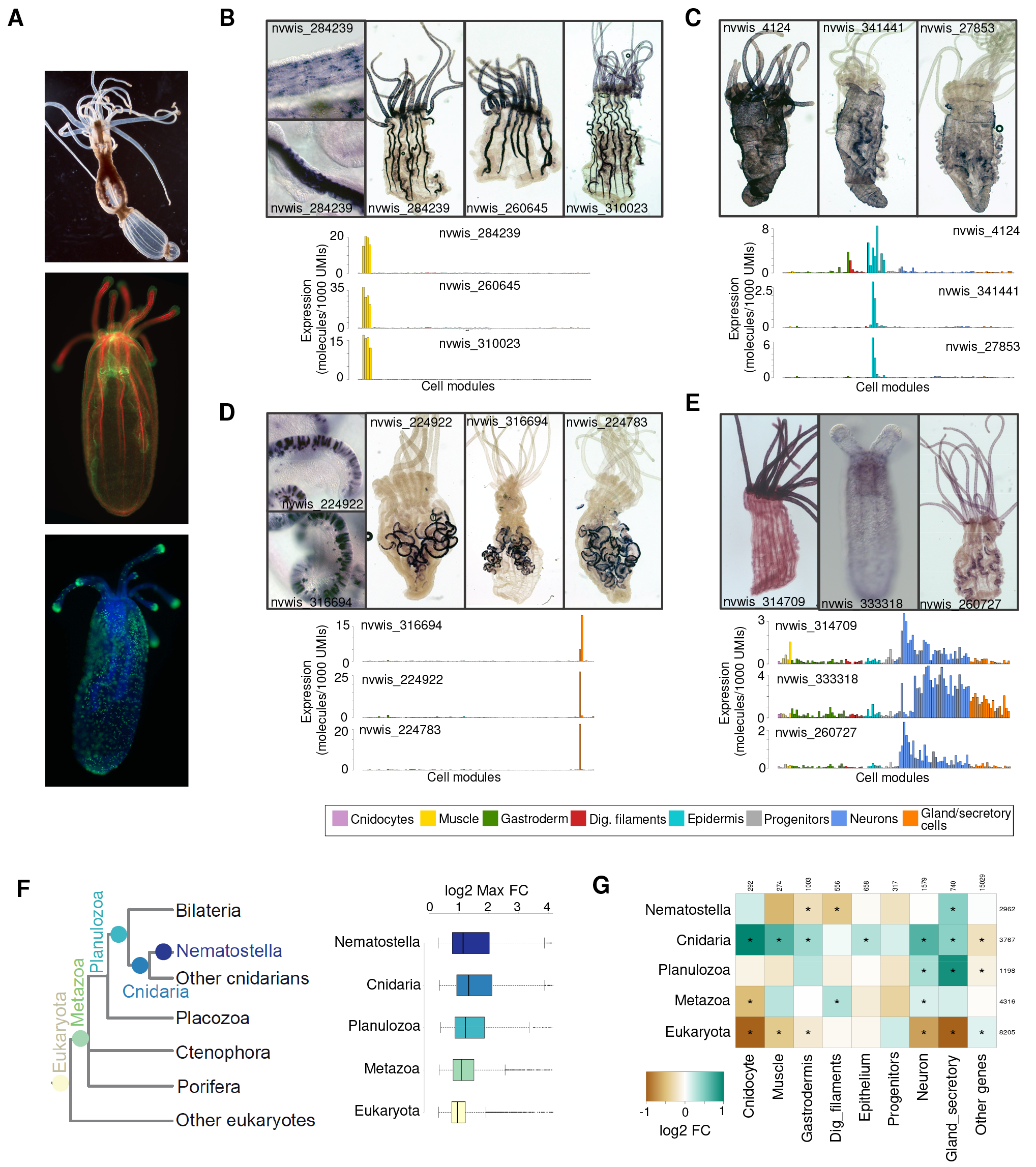
ISH validation of meta-cluster gene markers and phylogenetic distribution analyses. **(A)** Adult *Nematostella* polyp, oral opening and tentacles facing up (top). Juvenile polyp stained with phallodin (red), which binds to F-actin, and with an antibody against acetylated tubulin (green) (middle). Juvenile polyp stained with DAPI revealing nuclei (blue) and nematocysts (green) (green) (bottom). **(B)** Whole mount *in situ* hybridization in *Nematostella* polyps for three genes associated with muscle cell cluster. Barplots below represent the expression profile across the cell modules. **(C) (D) (E)** Same as (B), for genes associated with epidermal, gland and neuronal clusters, respectively. All expression values represented as molecules per 1,000 UMIs. **(F)** Gene expression variability across cell modules stratified by gene age. **(G)** Gene age enrichment/depletion in gene sets specific to each broad cluster. Asterisks indicate q-value < 0.001, chi-square test (BH correction).

Genes that are expressed broadly across tissues and developmental stages were shown to have older phylogenetic origins and are found across more diverse groups of species, while genes expressed in a narrower subset of tissues tend to have more recent phylogenetic origins (Sebé-Pedrós et al., 2016a; Tautz and Domazet-Lošo, 2011). To test if the same effect is observed for the detailed *Nematostella* transcriptional map, we annotated each predicted *Nematostella* protein to one of five phylogenetic classes, representing a spectrum of evolutionary depth ranging from the entire eukaryotic tree to proteins specific to *Nematostella* (**Figure 2F**). We next estimated the specificity of each gene in the dataset by computing the maximal fold change in expression levels across the cell modules we identified. As expected, we found that deeply conserved old genes generally show low cell type specificity, while new genes coding for Cnidaria (as defined by our current taxon sampling) are strongly enriched for tissue-specific expression (**Figure 2F** p≪0.001, Wilcoxon rank-sum). Complementarily, we next screened for gene age enrichments among the genes that are expressed specifically in each of the eight broad cell clusters described above. In agreement with the previous general observation of low cell type-specificity of paneukaryotic genes, we found that the depletion in tissue-specific ancient eukaryotic genes is common across broad cell clusters. In contrast, cnidocytes, a specific cnidarian cell type, are enriched in genes that originated at the stem of Cnidaria, linking gene innovation to the origin of new cell types (**Figure 2G**). Interestingly, a similar pattern is observed in muscle cell genes and this observation is in line with recent reports that suggested independent evolutionary origins of at least some muscle cell types in cnidarians and bilaterians (Steinmetz et al., 2012). Finally, neurons are significantly enriched in genes originated at multiple evolutionary times within Metazoa, indicating a step-wise assembly and specialization of the neuronal toolkit, from ancestral pan-metazoan genes to lineages-specific gene acquisitions (Liebeskind et al., 2016).

The detailed map of transcriptional states in *Nematostella* opens the way to analysis of the regulatory mechanisms underlying coordinated gene modules within putative transcriptional states and cell types. To test if the observed transcriptional programs are accompanied by expression of a comparably rich transcription factor (TF) repertoire, we identified 409 TFs expressed in our single cell cohort (with at least 20 total detected molecules). Of these TFs, 51% (207) showed evidence for module-specific expression (compared to 35% overall for all genes, p≪0.001, chi-square). Overall, we found that a median of 1.7% of the total transcripts in cell modules represent TFs (**Figure 3A**). This rate of TF expression was however tissue-specific, with 2-fold less TF expression in cnidocytes, muscle and gastroderm compared to neuronal and progenitor cell modules (p ≪ 0.001, chi-square). Classification of the 409 expressed *Nematostella* TFs into 38 structural families (**Figure 3B**) showed that homeobox TF class constitutes 26% of the total TFs expressed in *Nematostella*. Moreover, analysis of evolutionary conservation showed that 85% of the identified TFs are metazoan innovations (Wallberg et al., 2004), with candidates for cnidarian or *Nematostella* novelties represented in all structural classes, but enriched for C2H2 zinc fingers (p≪0.0001, chi-square). These results emphasize the relevance of metazoan TF innovations in animal cell type function, and in particular of the metazoan-expanded homeobox TF class (de Mendoza et al., 2013).

**Figure 3.**
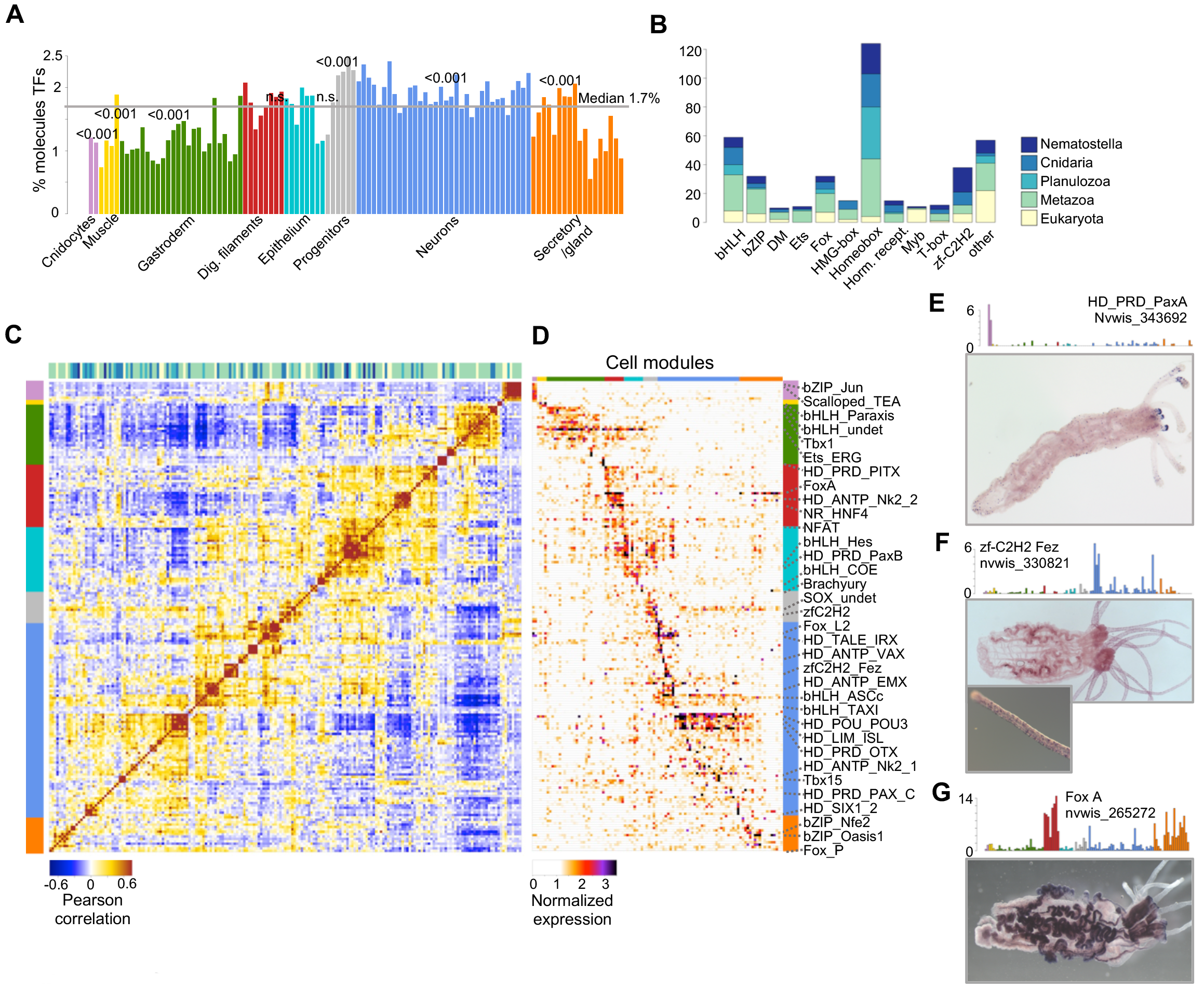
Transcription factor regulatory programs in *Nematostella*. **(A)** Structural class and evolutionary age of 409 detected *Nematostella* TFs. **(B)** Proportion of TF RNA molecules in each cell module. Chi-square test p-values for each broad cluster are indicated. **(C)** TF-TF Pearson correlation based on expression profile across cell modules, for TFs with >20 total molecules and FC>2 in at least one cell module. **(D)** TF expression profile across cell modules. **(E)** Expression profile and whole-mount ISH of *Nematostella* PaxA TF. **(F)** Expression profile and whole-mount ISH of *Nematostella* zfC2H2_Fez TF. **(G)** Expression profile and whole- mount ISH of *Nematostella* FoxA TF. All expression values represented in barplots as molecules per 10,000 UMIs.

Clustering of the TF-TF correlations among 207 TFs with module-specific expression (**Figure 3C**) revealed a remarkably rich combinatorial structure, identifying 24 groups of correlated TFs consisting of 2-21 TFs each. Analysis of the TF expression profiles across cell modules (**Figure 3D**) highlighted several TFs that are widely expressed within one of the eight broad clusters identified above, as well as many TFs with more specialized profiles. The detection of such broadly expressed TFs in the cnidocytes, muscle cells, digestive filaments, epidermis and neurons suggests the existence of tissue-specific hierarchy in *Nematostella* gene regulation in addition to the finer-grained and likely cell type-specific transcriptional signatures. We used ISH to morphologically cross-validate some of these observations of restricted, cell type-specific TF expression. For example, the cnidocyte specific PaxA expression observed in the scRNA data was shown to correlate by ISH with cnidocytes in the epidermis and tentacle tips (**Figure 3E**). In the case of the zf-C2H2 Fez TF, it showed expression in a specific population of neurons (**Figure 3F**); while the bilaterian foregut marker FoxA, which has been previously described to be broadly expressed in the digestive filaments and pharynx (Martindale et al., 2004), was expressed across digestive filament and gland cell clusters (**Figure 3G**). Overall, we uncover the TFs underlying specific states and cell type transcriptional identities in *Nematostella,* showing the potential of this approach to reconstruct the evolution of TF regulatory programs.

TFs drive tissue-specific transcriptional regulation by engaging regulatory elements in gene promoters and enhancers, through sequence-specific TF-DNA interactions. Since the sequence specificities of *Nematostella* TFs are not characterized, we sought to use predicted affinity models from protein sequence homologies, obtaining predicted binding motifs for 255 TFs (Weirauch et al., 2014) (**Figure S4A**). We then defined computationally 32,296 gene promoters in *Nematostella*, and extracted the sequence elements (-200/+50bp around annotated transcription start sites) associated to the genes specifically upregulated in each of the 104 cell modules introduced above. Statistical analysis detected enrichment (FDR<0.02) for specific binding motifs in all of the gene sets (**Figure 4A**). Furthermore, in-depth analysis identified cases in which the transcriptional profile of a TF was correlated with the enrichment profile of its putative binding motif, as exemplified by the FoxA-gland cells/digestive filaments profile, the ASC neuronal profile, ERG gastrodermal profile, and Pou neuronal/cnidocyte profile (**Figure 4B-E**). Spatial analysis showed that the enrichment of specific binding sites for FoxA, ASC, ERG and Pou is focused to the 200 bp upstream the annotated gene TSS. Analysis of shuffled controls (**Figure S4B**) confirmed the specificity of this result.

**Figure 4.**
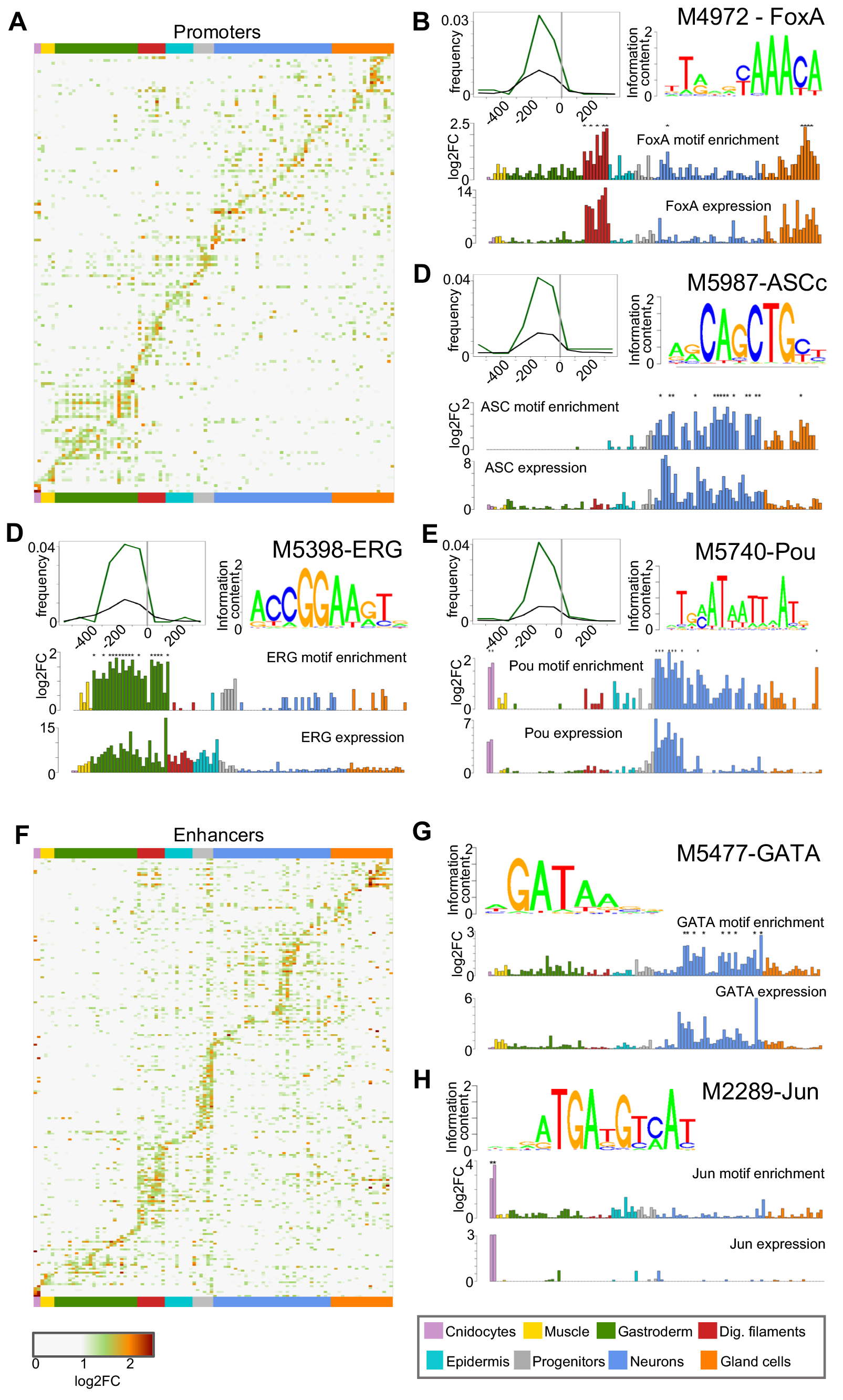
Motif enrichment analysis in *Nematostella* regulatory elements. **(A)** Heatmap showing significant (FDR < 0.01) specific motif (rows) enrichment in the promoters of cell module-specific gene sets (columns). **(B)** FoxA TF motif fold-change enrichment in 100bp windows around the TSS for a particular gene set with a significant motif enrichment (green line, FDR < 0.01) and the TSS of background genes (black line) (top-left). Motif logo derived from promoters with significant FoxA motif (top-right). Promoter motif enrichment (log2 Fold change) across cell modules (middle). FoxA expression (molecules per 10,000 UMIs) profile across cell modules (bottom). **(C) (D) (E)** same as (B) for ASCc, ERG and Pou TFs, respectively. **(F)** Heatmap showing significant (FDR < 0.01) motif enrichment in the enhancers associated to cell module-specific gene sets. **(G) (H),** Same as (B) for GATA and Jun TFs in enhancers. Asterisks indicate significant enrichments (FDR < 0.001). IDs represent CisBP entries (Weirauch et al., 2014).

Distally located enhancers play an essential role in tissue-specific gene expression in vertebrates and invertebrates (Dunham et al., 2012; Ho et al., 2014; Javierre et al., 2016; Kvon et al., 2014; Thurman et al., 2012; Vierstra et al., 2014). To examine if this is the case in *Nematostella*, we performed motif sequence enrichment analysis on 5,747 potential enhancer elements (Schwaiger et al., 2014), grouping them by proximity to the genes specifically upregulated in each of the cell modules. This gave rise to another set of putative links between TFs and tissue-specific regulation (**Figure 4F**), as exemplified by GATA and Jun TFs, associated to neuronal and cnidocyte cell clusters, respectively (**Figure 4G-H**). Importantly, this association was distinct from the preferences associated with the promoter sets (**Figure S4C**). Together, this analysis indicates that TF binding preferences are sufficiently conserved to allow modelling and association of sequence specificities with putative regulatory mechanisms. Moreover, the complex regulatory lexicon found at *Nematostella* enhancer elements indicates extensive distal regulatory element usage associated to cell type-specific transcriptional regulation in this early-branching metazoan, a feature that has been suggested to be an important novelty linked to the emergence of animal multicellularity (Sebé-Pedrós et al., 2016b).

The richness of the single cell RNA-seq profile (**Figure 1G**) allows us to examine transcriptional programs and possible cell subtypes at high resolution within larger cell clusters. In the case of the broad and heterogeneous neuronal cluster identified above, our data was consistent with previous studies that identified the proteins ELAV and ASC as important regulators of neural identity in *Nematostella* (Layden et al., 2012; Nakanishi et al., 2012) (**Figure S2**). In addition, our analysis adds a large number of neuronal-associated genes in *Nematostella,* including members of multiple ion channel families, synapsin, and peptide processing enzymes (**Figures 5A** **and S2**). In order to improve the resolution of neuronal cell maps, we set out to enrich this neuronal population by targeted isolation and sequencing of neurons. To this end, we used two existing *Nematostella* reporter transgenic lines driving expression of the fluorescent mOrange gene under the control of the ELAV promoter (Nakanishi et al., 2012) (**Figures S5A, S5B and S5E**) or the SoxB2a promoter (Richards and Rentzsch, 2014) (**Figures S5C, S5D and S5F**), two genes associated to neuronal cells. We sorted and processed 1,500 mOrange+ adult cells from each of these lines. Using the expression of the reporter gene itself, we could filter *a posteriori* negative cells, showing excellent agreement between Elav expressing cells and mOrange expressing cells (**Figure S5A**), the vast majority of which appear to be neurons. In contrast, in the case of the SoxB2a transgenic line, SoxB2a expression in neurons is not fully recapitulated by the reporter gene, possibly due to the genomic context of the reporter (**Figures S5C and S5G-H**).

**Figure 5.**
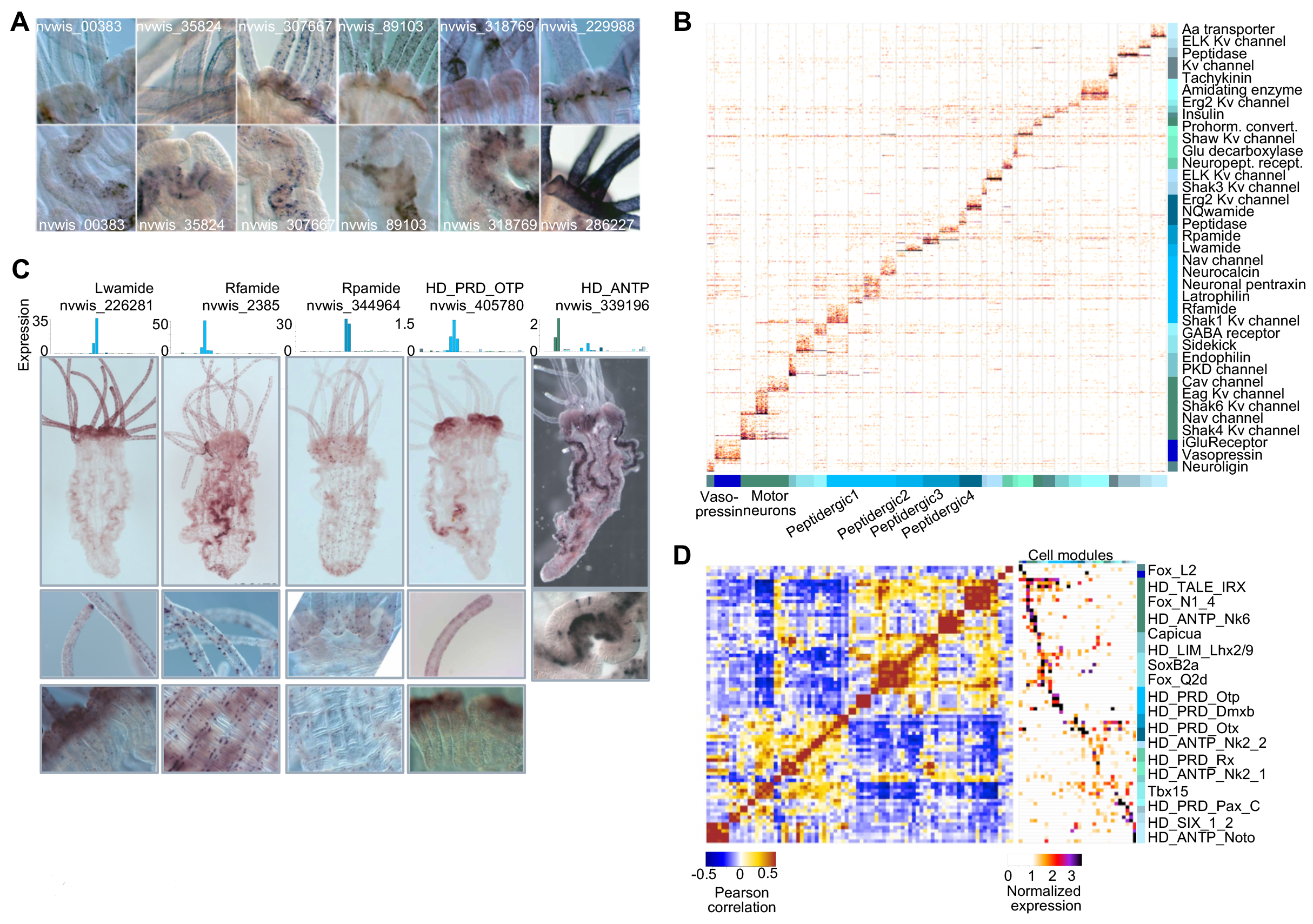
*Nematostella* neuronal cell cluster characterization. **(A)** Whole-mount ISH of genes associated to neuronal cell modules. **(B)** Normalized expression of 946 genes (rows) across 2,970 cells (columns), sorted by cell module association. For each module, the top 50 genes sorted by fold change versus the other modules were selected (with a FC threshold>=2). **(C)** Expression profile and whole-mount ISH of *Nematostella* Rfamide, Lwamide and Rpmide neuropeptides, and Otp and Nk6 TFs. All expression values represented as molecules per 1,000 UMIs. **(D)** Neuronal TF-TF expression correlation (left) and TF expression profile across cell modules (right).

We then zoomed in 4,775 neuronal cells, and constructed cell modules using 536 marker genes selected specifically as variable features within neurons. We used bootstrap analysis on the derived modules to select 32 robust neuronal cell modules and define genes and specific TFs linked with them (**Figures 5B-D**). Using known neuronal function-associated genes, we could re-define the *Nematostella* neuronal cell types in a systematic fashion (**Figures 5B**). A first subset of modules showed the characteristics of peptidergic neurons, showing specific expression of cnidarian neuropeptides RFamide, RPamide, LWamide and NQWamide (**Figures 5B-C**). ISH examination of some of these patterns in adult polyps revealed widespread expression in multiple spatially distinct territories and specific association with transcription factor profiles, such as the co-expression of RFamide with the homeodomain transcription factor Otp (**Figure 5C-D**). A second group of cells consisted of putative motor neurons (M), expressing specific TFs (**Figure 5C**) and a high number of sodium, calcium and potassium ion channels (20 different channels expressed in these modules compared with a median of 3 in the other neuronal clusters; p≪0.0001, chi-square). Finally, we identified multiple additional modules with highly specific expression profiles and unique combinations of transcription factors but of unknown functional characterization. Our results suggest that cnidarians host a greater diversity of neuronal cell types than previously acknowledged, and that they share transcriptional parallels with bilaterian nervous systems. Future similar analyses in additional species will reveal the extent to which cnidarian and bilaterian nervous systems share neural features, and they will help to trace lineage-specific innovations of neuronal cell types.

We performed a similar in-depth analysis for cnidocyte and gland/secretory cells (**Figure S6**). We found that cnidocytes form three distinctive groups (Zenkert et al., 2011) that likely correspond to mastigophores (expressing spinalin, NEP proteins, and minicollagen 1), haplonemas (weakly expressing mastigophore markers as well as progenitor cell markers such as SoxB2a (Richards and Rentzsch, 2014)), and spirocytes (expressing two specific minicollagens) (**Figure S6A-C**). On the other hand, re-analysis of the extremely varied cells within the gland/secretory broad cluster revealed multiple cell types expressing unique combinations of digestive enzymes and venom proteins (**Figure S6D-F**).

In this work, we decomposed transcriptional landscapes at single cell resolution in the non-bilaterian animal *Nematostella*. The map of cell types we derive re-organizes *Nematostella* cells into eight broad categories, and uncover unanticipated diversity within them. Remarkably, this complex cell type hierarchy is supported by highly specific co-expression of a large TF repertoire, and by a corresponding diverse motif lexicon encoded in promoters and enhancers associated to cell type-specific genes. The detection of this rich pool of tissue-specific TFs and motif signatures in *Nematostella* suggests that highly-organized tissue-specific gene expression programs are not an exclusive hallmark of complex metazoans but may have already existed in the cnidarian-bilaterian ancestor. However, it remains to be determined if and to which extend cnidarians implement those programs in a similar way than vertebrates or other bilaterian models (Dunham et al., 2012; Ho et al., 2014; Sexton et al., 2012), in particular regarding the balance between direct regulation of proximal promoters by sequence-specific factors versus indirect, epigenetically-mediated, control of repressive and active genomic domains (Bonev and Cavalli, 2016).

The methodology we employ here aims at unbiased sampling of cells from a whole organism. Our approach may under-sample or miss completely very rare cell types, or cells that are particularly sensitive to the sorting process. However, the richness of the derived maps, and the possibility to sample cells directly from specimens without the need for culturing or complex preparation, suggest that single cell analysis may quickly become a method of choice for studying gene regulation in non-model organisms. Moreover, while our analysis used the existing *Nematostella* reference genome and its annotation, we envision applications of this methodology to more poorly characterized species and genomes, opening the way to massive expansion in the phylogenetic coverage and overall quality of models for genome regulation. Such future systematic studies may provide unexpected new insights into the evolution of cell types and the associated genome regulatory programs in metazoans.

## Acknowledgements

We thank David Lara-Astiaso for critical comments on the manuscript, Manuel Irimia for discussion on cross-species clustering, and all the members of the Tanay lab for comments and discussion. We thank Dr. Malte Paulsen and the EMBL flow cytometry facility for assistance with initial cell sorting trials. We thank Dr. Fabian Rentzsch for providing the Elav and SoxB2a reporter lines and Dr. Detlev Arendt (EMBL) for allowing us to use riboprobes that were generated by H.M. during time spent in his laboratory, as well as for constructive discussions regarding the definition and evolution of cell types. A.S.-P. is supported by an EMBO Long-Term Fellowship (ALTF 841-2014). Research in H.M./F.S. group was supported by the Region Ile de France (program SESAME 2016 “Paris Single Cell Centre”) and Pasteur Citech (“Single cell genomics”). Research in A.T. group was supported by the European Research Council. A.T. is a Kimmel investigator.

## Methods

### *Nematostella* culture, dissociation and sorting

*Nematostella* polyps were spawned and reared as previously described to 11 days (“tentacle bud stage”), and five months (small adult polyps) (Fritzenwanker and Technau, 2002; Hand and Uhlinger, 1992; Stefanik et al., 2013). Multiple polyps (~20) from the same life stages (adult or juvenile) where processed together, resulting in a unique single cell suspension from where cells were randomly sampled by FACS sorting, as described below. This process was repeated successively in order to ensure a short time between dissociation and cell capture (<3h), eventhough cell viability was continuously monitored throughout the sorting. The dissociation and sorting were done using with the same reagents (enzymes, fluorescent sorting labels, and media), in order to minimize technical factors and various batch effects.

Cells were distributed into 384-wells capture plates (all coming from the same production batch) containing 2 ul of lysis solution using a using a FACSARIA III cell sorter. Lysis solution contain 0.2% Triton and RNAse inhibitors plus barcoded poly(T) reverse-transcription (RT) primers for single cell RNA-seq. Doublet/multiplet exclusion was performed using FSC-W vs FSC-H. Additionally, in the case of ELAV and SoxB2 reporter transgenic lines, we selected mOrange-positive cells. Fresh cell dissociates were prepared every 3h and sorted plates were immediately spun down, to ensure cell immersion into the lysis solution, and frozen at −80°C until further processing. Eight empty wells were kept in each plate as a control for data analysis.

### Massively Parallel Single-Cell RNA-seq (MARS-seq)

Single cell libraries were prepared as previously described (Jaitin et al., 2014). All 17,664 single cell libraries (starting from 46 384-wells MARS-seq capture plates) were prepared with the same conditions (incubation times, temperatures, etc) and reagents. First, Using a Bravo automated liquid handling platform (Agilent), mRNA was converted into cDNA with an oligonucleotide containing both the unique molecule identifiers (UMIs) and cell barcodes. Unused oligonucleotides were removed by Exonuclease I treatment. cDNAs were pooled (each pool representing half of the original 46 384-wells MARS-seq plate, 92 batches in total) and linearly amplified using T7 in-vitro transcription and the resulting RNA was fragmented and ligated to an oligo containing the pool barcode and Illumina sequences, using T4 ssDNA:RNA ligase. Finally, RNA was reverse transcribed into DNA and PCR amplified. Resulting libraries were tested for amplification using qPCR and the size distribution and concentration were calculated using Tapestation (Agilent) and Qubit (Invitrogen). scRNAseq libraries were pooled at equimolar concentration and sequenced using Illumina NextSeq 500 sequencer, in three sequencing runs, using high-output 75 cycles v2 kits (Illumina). We obtained 923M reads, of which 84% passed filtering, resulting in 785M reads with an average depth of 38,000 reads per cell and 6 reads/UMI on average.

### Whole-mount *In Situ* Hybridization and Imaging

Fixation of tentacle bud and adult stage *Nematostella* was carried out as previously described (Wolenski et al., 2013). *In situ* hybridization with digoxigenin-labeled riboprobes at a final concentration of 1 ng/uL was performed as previously described. Following *in situ* hybridization, polyps were sequentially cleared through a 40/60/70% glycerol series and imaged on a Zeiss AxioImager DIC microscope or a Zeiss Discovery V16 microscope. Color balance, contrast and levels were adjust across the image and images were cropped using Adobe Photoshop.

For *in situ* probes transcribed from PCR based templates, the primer sequences can be found in the table below. For *in situ* probes generated from templates produced via standard cloning and restriction digest.

### Staining of nematocyte capsules, tubulin and actin

Polyp fixation and staining were done as previously reported (Marlow et al., 2009). Briefly, polyps were fixed in 4% paraformaldehyde and 0.2% gluteraldehyde for 90 seconds, followed by 4% paraformaldehyde for one hour at 4C, and then washed five times into PBS (18.6 mM NaH2PO4.H20, 84.1 mM Na2HPO4.2H2O, 1.750 mM NaCl). Polyps were blocked for 30 min at room temperature and incubated overnight at 4C in mouse anti-acetylated tubulin primary antibody (Sigma, T6793) and then washed out three times into PBS-triton. Polyps were subsequently incubated in Alexa Fluor 488 goat anti-mouse secondary antibody (Fisher Scientific, 10256302) and post-stained with Phalloidin (Fisher Scientific, 10656163). Polyps stained for nematocyst capsules were fixed and treated according to Marlow et al. 2009 (Marlow et al., 2009) and following Szczepanek, el. Al. 2001 (Szczepanek et al., 2002). Basically, if calcium chelators are present during fixation, cnidocyte capsule walls can be stained with DAPI and visualized using a green filter under fluorescent illumination while nuclei (DAPI-stained DNA) are visible using a blue filter.

### Pre-processing and filtering of MARS-seq reads

Reads were mapped into *Nematostella* genome v1.0 (http://genome.jgi.doe.gov/Nemve1/Nemve1.home.html) using bowtie2 with default parameters and associated with a gene intervals defined by merging two existing *Nematostella* annotation(Putnam et al., 2007; Schwaiger et al., 2014). After merging, we extended gene intervals up to 2kb downstream or until the next gene in frame is found. This accounts for the poor 3′UTR annotation of *Nematostella* genome, which causes most of the MARS-seq (a 3′ biased RNA-seq method) reads to map outside genes (**Figure S1B**). Mapped reads were further processed and filtered as previously described (Jaitin et al., 2014). UMI filtering include two components, one eliminating spurious UMIs resulting from synthesis and sequencing errors, and the other eliminating artifacts involving unlikely IVT product distributions that are likely a consequence of second strand synthesis or IVT errors. The minim FDR q-value required for filtering was 0.2. We filtered cells with less than 100 UMI from downstream analysis.

### Gene selection and cell module detection

We used the MetaCell package to select gene features, construct gene modules and create projected visualization of the data, using parameters as described below. We applied preliminary cell filtering based on total UMI counts using a permissive threshold of 100 UMIs. For gene selection we used a normalized depth scaling correlation threshold T_gr_ of −0.05. For module cover construction we used K=250, minimum module size of 50 and automatic filtering of background noise using an initial epsilon value of 0.03. Bootstrapping was performed using 1000 iterations of resampling 75% of the cells leading to estimation of co-clustering between all pairs of single cells, and identification of robust clusters based on single or grouped modules. For 2D projections, we used a K-nn constant of 80, and restricted the module graph degree by at most 8. These parameters were used for both global clustering (**Figure 1**) and clustering of subsets of cells, with only module cover construction K-nn parameter and minimum module size being adjusted in each case for in-depth analysis of neuronal cells (K=150, min size=30), cnidocytes (K=120, min size=20) and gland/secretory cells (K=150,min size=30).

We performed manual validation and adjustment of the automatic module covers in Figure 1 as follows. We eliminated a total of 12% of the cells that were part of modules with low self-consistency (less than 20% of the edges in the K-nn graphs involving at least one cell in a module connecting two members of the module). We also filtered modules that were not enriched by at least one gene at over 9 fold over the median of the entire populations. We then grouped and labelled modules by combining gene annotation and results from bootstrap analysis, organizing all cells into 8 broad groups and supporting their robustness by testing the association of each cell to the broad cluster across bootstrap sampling iterations (**Figure 1H**).

In order to deriving clusters (rather than modules) and supporting their robustness in the analysis of neuronal transcriptional states in Figure 5 and Figure S9, we used both the bootstrap robustness analysis and a parametric test for module specific transcriptional enrichment. First, groups of cell modules with >35% shared cells co-clustering in our 1000 bootstrap replicates were merged. Following such merging, modules for which the average probability of co-clustering across all pairs in 1000 bootstrap iterations were less than 70% filtered. Minimal enrichment was then tested for each merged module by identifying a set of module-specific genes (top 50 genes with FC>=2) and computing the top 1% of their total expression across all non-module cells (excluding also cells in the two most similar modules). Given this top percentile as a threshold, the fraction of cells in the module that express the module's genes over the threshold was computed, and additional filtering was applied if this value was lower than 50%. We note that cells that were filtered during this combined scheme may be part of additional undetected states, or may represent weaker signal that is in fact part of other, more robust modules. Nevertheless, for the purpose of our analysis here, the main goal was to ensure the robustness of the clusters that can be supported by the data unambiguously, and it must be assumed that additional rare or weaker behaviour are left uncharacterized.

### Gene functional annotation

We used blastp (with parameters *–evalue* 1e-5 and *-max_target_seqs* 1) to find for each protein of the merged *Nematostella* predicted proteome (JGI plus Vienna annotation) the most similar, if any, human, drosophila and yeast homologs (retrieved from Uniprot). Additionally, we predicted for each protein the Pfam domain composition using Pfamscan (Punta et al., 2012) with default curated gathering threshold. *Nematostella* TFs were identified using univocal Pfam domains for each structural TF family (de Mendoza et al., 2013). In the case of multi-TF families (Homeobox, Fox, bHLH, bZIP, DM, Smad, Myb, NR, RFX, RHD, SRF, Ets, T-box and Sox), we used phylogenetic analyses for each family in order to classify them into specific subfamilies (together with the complete TF sets of additional 10 animal species, including *Homo sapiens* and *Drosophila melanogaster* for reference annotation). Briefly, sequences were aligned using MAFFT (Katoh et al., 2002), the resulting analysis were manually edited, ProtTest (Darriba et al., 2011) was used to define the best-fit aminoacidic substitution model in each case, and then phylogenies were computed using RAxML (Stamatakis, 2006) and Phylobayes (Lartillot et al., 2009), for maximum likelihood and Bayesian inference, respectively.

### Motif analysis

Inferred DNA-binding motif preferences for *Nematostella* TFs were obtained from CisBP database (http://cisbp.ccbr.utoronto.ca/) (Weirauch et al., 2014). Briefly, experimentally determined Position Weight Matrice (PWM) from different species are transferred to other species TF given a certain threshold of primary protein sequence identity. In the case of *Nematostella,* CisBP contains 1,347 PWMs (excluding Transfac) corresponding to 255 *Nematostella* TFs. In order to reduce redundancy, we computed the binding energy of these 1,347 PWMs in all *Nematostella* promoters (TSS −200/+50 bp) and calculated the motif-motif correlation based on these energies. A correlation threshold of 0.7 was used to cluster motifs and a single representative of each cluster was selected for downstream analyses, resulting in a reduced set of 384 PWMs (**Figure S4A**).

We extracted promoters sequences using −200:+50 bp from annotated TSSs and associated sequences with cell modules whenever their gene was at least two fold over-expressed in the module compared to the background. Enhancers sequences were defined based on Schwaiger et al. 2014 (Schwaiger et al., 2014) and grouped into cell modules if their closest TSS was induced in the module at least two fold. For a short sequence element **s[1..k] = s_1_,..,s_k_**, and a PWM **w_i_[c]**, the standard local probability model is defined by multiplication: **log(P(s)) =** \**sum_i_log(w_i_[s_i_])** and the binding energy for a larger sequence element can be approximated(Tanay, 2006) by **E(s[1..n]) = log(**\**sum_(j = 1:(n-k))_P(s[j:(j+k)])**. For each PWM, the 0.98 quantiles of genome-wide binding energies in windows of 250bp (for promoters) or 150bp (for enhancers) were determined. These quantiles values were then used as thresholds to determine motif occurrence for each PWM at each element. The enrichment level of each PWM/cell module pair was computed as the fold change between the frequency of occurrence of a motif in the cell module's promoters/enhancers and the frequency in the background gene set (all other genes detected in this study). Enrichments were assessed statistically using a hypergeometric test. We account for multiple testing by performing 100 random permutations of the promoter-motif energy matrix, computing p-values for each permutation and using the resulted distribution to derived FDR values on the empirical enrichments. An FDR threshold of 0.02 was used for the motif enrichment visualization. Additionally, only motifs with a fold change enrichment over 1.5 in at least one cell module, and a minimum foreground count of 5 (ie. at least five genes in the cell module gene set with the motif in their promoters/enhancers) and a background count of 100 in this module were considered.

### Phylogenetic distribution estimation

We used the complete predicted proteomes of 31 species at key phylogenetic positions in order to compute orthogroups, including an extensive set of 10 cnidarian species (*Acropora digitifera, Aiptasia pallido, Anthopleura elegantissima, Edwardsiella lineata, Fungia scutaria, Nephthyigorgia* sp., *Alatina alata, Atolla vanhoeffeni, Hydra magnipapillata, Podocoryne carnea*) (Babonis et al., 2016), 6 other planulozoans (*Homo sapiens, Branchiostoma floridae, Drosophila melanogaster, Tribolium castaneum, Capitella teleta, Lottia gigantea*), 6 other metazoans *(Trichoplax adhaerens*, *Amphimedon queenslandica, Oscarella carmela, Sycon ciliatum, Mnemiopsis leidyi, Pleurobrachia bachei*) and 8 non-metazoan eukaryotes (*Salpingoca rosetta, Capsaspora owczarzaki, Creolimax fragrantissima, Saccharomyces cerevisiae, Spizellomyces punctatus, Dictyostelium discoideum, Arabidopsis thaliana, Naegleria gruberi*). We computed reciprocal blast results between all complete proteomes, with fixed database size and e-value threshold of 1e-04. Based on these reciprocal blast results, orthogroups were computed using orthoMCL algorithm (Li et al., 2003) with an inflation value (*I* parameter) of 1.3. We parsed these orthogroups using a parsimony criterion in order to generate an age estimation for each *Nematostella* gene.

### Cross-species transcriptome comparison

We used the normalized expression values for amniote organ-specific transcriptomes generated by Brawand *et al*. (Brawand et al., 2011). *Nematostella*-human orthologs (see above) were used to merge this matrix with *Nematostella* meta-cluster gene expression (represented by fraction of total molecules in the meta-cluster). Both matrices were centered before merging and then quantile normalized. We then used the normalized expression profile of 1,253 highly variable genes across samples (with a fold-change > 2 in at least 4 organ samples and a fold-change > 2 in at least 1 *Nematostella* meta-cluster) to compute the Pearson correlation between samples.

---

**Figure S1 *Nematostella* scRNA-seq UMI statistics and cell module analysis. (A)** Fraction of cells in each cell module originating from each of the batches (each batch representing half a 384-wells MARS-seq plate, see Methods). **(B)** Distribution of mapped reads around Transcription End Sites (TES) of *Nematostella* originally annotated genes, we expanded the 3’ end of each gene 2Kb (or up to the next gene in the same strand) to capture reads mapping downstream of the original TES. **(C)** Correlation of total UMIs per cell for all genes (x-axis) versus total UMIs per cell for the top-5 size correlated genes. **(D)** Cumulative distribution of number of marker genes detected per single cell. **(E)** Total number of cells per module. **(F)** Total number of molecules per module. **(G)** Clustering robustness analysis. The heatmap shows the Pearson correlation (based on the expression profiles of the 300 marker genes) between two clustering solutions based on random selection of half of the cells. Color bars identitate the new modules based on similarity to the original clustering broad clusters (see **Figure 1**). **(H)** Bootstrap analysis. Heatmap representing the frequency of cell-to-cell association in 1,000 bootstrap subsamplings. **(I)** Number of genes per cell module with a fold change >=2.

**Figure S2. Gene expression profiles.** Expression profiles of selected genes in each cell module. All expression values are shown as molecules per 1,000 UMIs.

**Figure S3. Cross-species tissue transcriptome comparison. (A)** Pearson correlation between organs/tissues (color-coded in the first colorbar) of different animal species (color-coded in the second colorbar). **(B)** Normalized expression of the gene orthologues (rows) used to compute the organ/tissue (columns) correlations. Vertebrate organ transcriptome data obtained from (Brawand et al., 2011).

**Figure S4. *Nematostella* motif enrichment supplementary analyses. (A)** Motif-motif Pearson correlation based on occurrence of each motif in *Nematostella* promoters. A single representative PWM from each cell module (indicated in black in the flanking colorbars) was used for downstream analyses. The PWMs were obtained from CisBP database (Weirauch et al., 2014). **(B)** Cumulative distribution of p-values for the hypergeometric motif enrichment tests in promoters and enhancers in the original dataset (black) and shuffling cell module-gene associations (red). **(C)** Motif-motif Pearson correlation between shared promoter (columns) and enhancer (rows) enriched motifs (presented in the same order as promoter motifs in Figure 4), showing that there is no correspondence of the same motif enrichments at enhancers and promoters.

**Figure S5. *Nematostella* ELAV and SoxB2a reporter lines scRNAseq analysis. (A)** Expression of mOrange reporter and ELAV genes across the 17 modules defined from 1,500 sorted Elav+ cells. **(B)** Heatmap showing the Pearson correlation (based on the expression profiles of the 413 marker genes used to cluster Elav+ cells) between Elav+ cell modules (columns) and the general modules (rows, see **Figure 1**). The expression of Elav in the general cell modules is shown as an horizontal barplot. Elav cell modules mapping to neuronal modules are highlighted with a blue rectangle. **(C) (D)**, Same as (A) (B) for 1,500 SoxB2a+ cells. **(E)** 2d projection of the Elav+ cells, neuronal cells are highlighted in blue. **(F)** same as e for SoxB2a+ cells. **(G)** Chromatin features around the ELAV gene. The promoter sequence cloned for the Elav+ reporter line is highlighted in red (Nakanishi et al., 2012). **(H)** Same as g for SoxB2a (Richards and Rentzsch, 2014). Notice that in the case of SoxB2a there are multiple regulatory elements bot upstream and downstream of the cloned promoter and whose regulatory inputs are not captured by the SoxB2a reporter line. This may explain the discrepancies observed between SoxB2 and reporter mOrange expression. All expression values are shown as molecules per 1,000 UMIs. Chromatin and bulk RNA-seq data from Schwaiger et al. (Schwaiger et al., 2014).

**Figure S6. *Nematostella* cnidocyte and gland/secretory broad clusters characterization. (A)** Normalized expression of 193 genes (rows) across 297 cnidocyte cells (columns), sorted by cell module association. For each module, the top 80 genes sorted by fold change versus the other modules were selected (with a FC threshold>=2). **(B)** Cnidocyte cells bootstrap analysis. Heatmap representing the frequency of cell-to-cell association in 1,000 bootstrap subsamplings. **(C)** Expression profiles of venom/spine proteins, TFs and progenitor markers across the cnidocyte cell modules. All expression values are shown as molecules per 1,000 UMIs. **(D)** Normalized expression of 283 genes (rows) across 297 gland/secretory cells (columns), sorted by cell module association. For each module, the top 100 genes sorted by fold change versus the other modules were selected (with a FC threshold>=2). **(E)** Gland/secretory cell modules bootstrap analysis. Heatmap representing the frequency of cell-to-cell association in 1,000 bootstrap subsamplings. **(F)** Expression profiles of selecter markers across gland/secretory cell modules. Proteins like phospholipases A2, astacins or CAP domain proteins have been described to be related to venom functions in cnidarians (Jaimes-Becerra et al., 2017; Moran et al., 2013). All expression values are shown as molecules per 1,000 UMIs.

